# A Male Expressed Copulatory Plug Increases Female Fertility in *Caenorhabditis* Nematodes

**DOI:** 10.64898/2026.06.29.735219

**Authors:** Lynsey J. Blevins, Andrew Grieve, Michael Sauria, Erik C. Andersen, Katja R. Kasimatis

**Affiliations:** Department of Biology, University of Virginia, Charlottesville, VA, 22904, USA; Department of Biology, Johns Hopkins University, Baltimore, MD, 21218, USA

**Keywords:** Mate plugging, Mating signal, Reproductive benefit, *Caenorhabditis* nematodes

## Abstract

The genomes of self-fertilizing hermaphrodites provide an opportunity to probe why some genes with sex-specific functions are lost while others are maintained. In the androdiecious species, *Caenorhabditis elegans*, the presence of mate plugging is polymorphic across genetic backgrounds. The production of a mating plug is controlled by a single gene, *plg-1*, which has male-specific expression. Given that males are present less than 1% of the time, this gene is likely lost if mate plugging is maintained through intrasexual selection in males as is often hypothesized. We capitalized on a panel of almost 2,000 *C. elegans* natural strains to show that *plg-1* is maintained in at least 61% of genetic backgrounds. Genetic diversity estimates suggest that the loss of *plg-1* is correlated with species globalization. We demonstrate in both *C. elegans* and the gonochoristic species, *C. remanei*, that the presence of a mating plug does not prevent or delay subsequent males from mating. Instead, females with a mating plug have an extended peak reproductive window and higher fecundity than females without a mating plug. Together, these results indicate that mate plugging is female beneficial. Our results support that mate plugging might not indicate male competition and can instead represent reproductive cooperation.

## INTRODUCTION

Evolutionary transitions in mating system offer an opportunity to link changes in genome content and function with reproductive traits that benefit one sexual function over another. The transition from outcrossing to self-fertilizing is recurrent across plant and animal taxa (Shimizu and Tsuchimatsu 2014). A hallmark of self-fertilization is a streamlining and shrinking of the hermaphrodite genome because of inbreeding depression, the purging of deleterious alleles, and severe population bottlenecks (Oyama et al. 2008; Wright et al. 2008; Thomas et al. 2012). Additionally, populations with increased rates of self-fertilization have reduced intragenomic conflict due to the increased efficacy of purifying selection (Wright et al. 2008). Although typically observed as a reduction in selfish genetic elements (Wright et al. 2008; Thomas et al. 2012), it can also lead to a reduction in sexual conflict. In self-fertilizing hermaphroditic species, sexual conflict arises as interlocus conflict between the genes underlying female and male functions (Abbott 2010). If there is greater opportunity for selection to act on one sexual function over the other, then sex-specific selection should maintain the genes of that sexual function. Genes of the other sexual function are expected to be lost due to purifying selection or genetic drift. Thus, self-fertilizing hermaphrodites are candidate species for testing why, how, and what classes of genes are lost under reproductive shifts.

*Caenorhabditis* nematodes are an excellent model for studying the genomics underlying mating system transitions. The genus contains three independent lineage transitions – *C. elegans*, *C. briggsae*, and *C. tropicalis* – to androdiecious mating systems comprised predominantly of self-fertilizing hermaphrodites and rare (on average less than 1%) males (Cutter et al. 2019). The hermaphroditic species genomes are smaller than their gonochoristic sister species (Yin et al. 2018; Noble et al. 2021; Teterina et al. 2023; Wang et al. 2026), which is driven by a loss of genes with male-specific function. For example, the *C. briggsae* genome has 23.5% fewer protein coding genes than its gonochoristic sister species *C. nigoni*, and the *C. nigoni* genes lacking orthologs in *C. briggsae* are disproportionately male-biased in expression (Yin et al. 2018). This loss of male-specific genes underlies the degradation of successful male mating behaviors (Chasnov et al. 2007; Garcia et al. 2007; Thomas et al. 2012). Together, these patterns suggest a repeated evolutionary history of male gene loss due to weaker stabilizing selection on male function coupled with reduced sexual selection and sexual conflict.

An interesting case study for testing these processes is the ancestral and largely conserved gene *plg-1*. This gene is expressed exclusively in the male vas deferens and produces a mucin protein that is deposited over the female/hermaphrodite vulva after sperm transfer (Barker 1994; Palopoli et al. 2008). Such mating plugs are common across insects, reptiles, and some mammals (Parker 1970; Schneider et al. 2016). Mating plugs display a diversity of forms ranging from solid and deposited within the vaginal canal to gelatinous and deposited over the vulval opening (as seen in nematodes) (Barker 1994; Dixson and Anderson 2002; Schneider et al. 2016). This range of forms is hypothesized to arise through multiple non-mutually exclusive evolutionary processes (Voss 1981). Under a paternity assurance hypothesis, mating plugs are hypothesized to increase a male’s reproductive success by impeding another male from attempting to mate or physically preventing another mating. This competitive exclusion leads to strong intrasexual selection and can have a neutral effect on female fecundity or can cause sexual conflict if male competition limits female choice and access to sperm (Eberhard 2015; Schneider et al. 2016). Alternatively, mating plugs can directly benefit female fecundity by acting as a nuptial gift (Perry and Rowe 2008; Schneider et al. 2016), increasing ejaculate retention (Avila et al. 2015; McDonough-Goldstein et al. 2022), or increasing female remating rate (Perry and Rowe 2008; Yun et al. 2024). Correspondingly, male fecundity would also increase. *Caenorhabditis* nematodes offer a unique opportunity to examine the potential interconnectedness of these different functions and how they relate to male-specific selection. For example, under the paternity assurance hypothesis alone, *plg-1* would be expected to be lost in hermaphroditic *Caenorhabditis* species either because of relaxed selection on any male functions or selection on hermaphrodites against mating with mate-plugging males. Conversely, the female benefit hypothesis would indicate that *plg-1* would be maintained in hermaphrodites if the fitness benefit provided by a mating plug was sufficient to overcome the rarity of males and potential loss through stochastic events. Here, *plg-1* would be favored in males because the plug increases male fecundity and in hermaphrodites because their sons will express *plg-1* and therefore have higher fecundity. Previous work in *C. remanei*, found no evidence for paternity assurance and nominal evidence for female benefit (Timmermeyer et al. 2010), suggesting that *plg-1* could be selectively maintained in hermaphroditic species, though relaxed constraint is more likely.

In *C. elegans*, *plg-1* is polymorphic across genetic backgrounds (*i.e.*, strains) with some strains maintaining a functional copy and others having a nonfunctional gene copy. The loss of function is driven by retrotransposon insertion of a *Cer1* element and its long terminal repeats (LTRs) into the third exon of *plg-1* {Palopoli.2008; Fig. 1A}. Here, we connect the polymorphic nature of *plg-1* in *C. elegans* with its reproductive fitness effect to better understand if this gene is being purged from the genome due to male-specific function or if instead it is being selectively maintained. First, we expand the genotyping of *plg-1* across *C. elegans* strains to better understand the phylogenetic context and evolutionary history of this locus. Then, we explicitly test the paternity assurance hypothesis in both *C. elegans* and *C. remanei* in a robust, repeatable manner. We support these data by quantifying hermaphrodite and female reproductive fitness to test the female benefit hypothesis. Together, our work indicates that even a small fitness benefit to hermaphrodites can maintain a male-specific gene when males are rare.

**Figure 1.**
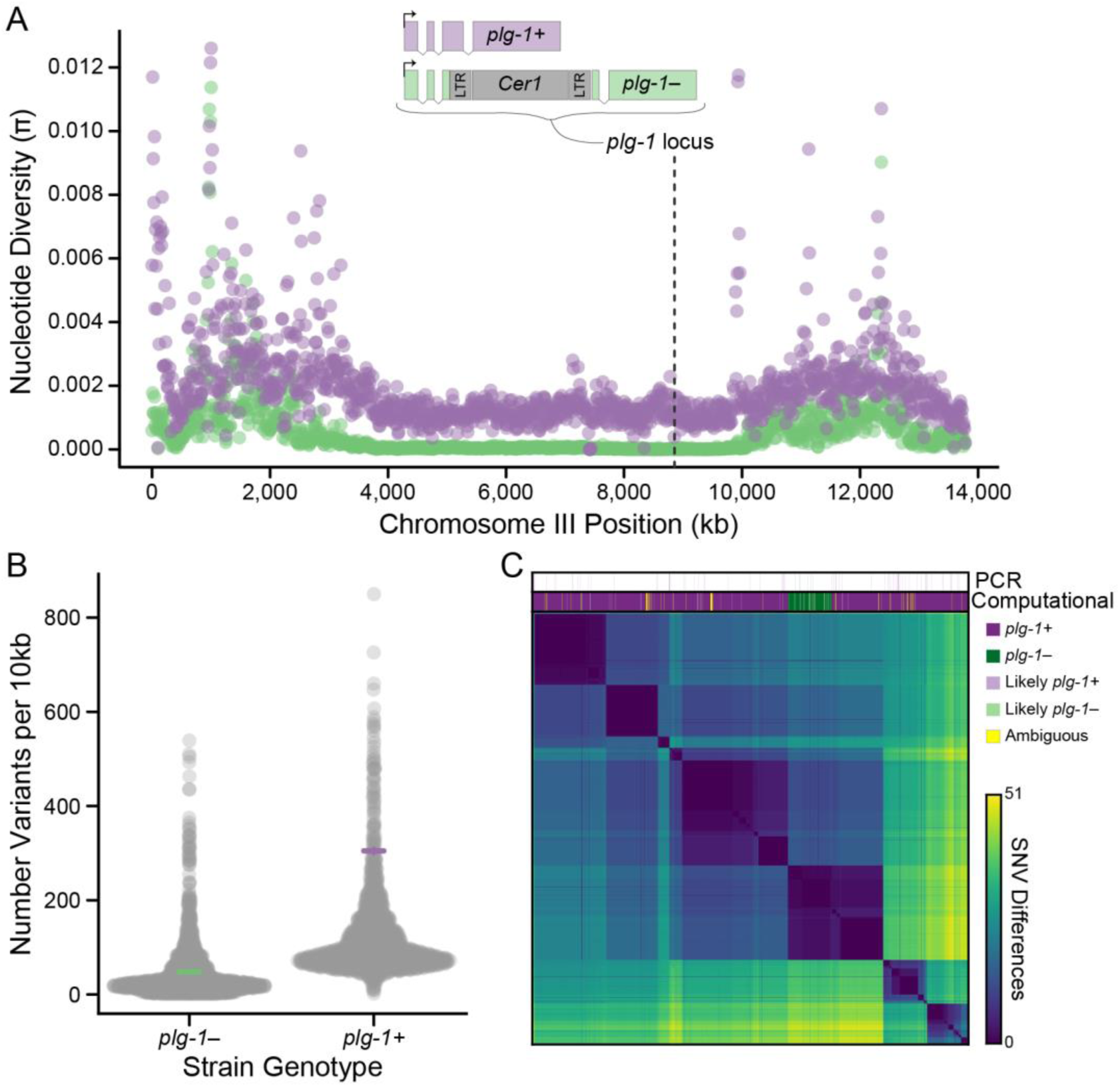
Chromosome III nucleotide diversity in 1,194 *plg-1+* and 187 *plg-1–* strains. **A)** Nucleotide diversity calculated in 10 kb sliding windows along chromosome III for *plg-1+* (purple) and *plg-1–* (green) strains. The dashed line shows the location of the *plg-1* locus with the inset representing the gene models of for functional and non-functional *plg-1* copies. Nucleotide diversity is highest on the chromosome arms in both genotype groups, though overall diversity is higher in *plg-1+* strains. **B)** The number of variants falling into each 10 kb sliding window on chromosome 3 for *plg-1+* and *plg-1–* strains. The mean and standard error for each genotype is given by the purple (*plg-1+*) and green (*plg-1–*) crosses. There are on average more variants per window in *plg-1+* strains. **C)** The pairwise difference in the number of single nucleotide variants (SNV) across all strain pairs. The computationally determined *plg-1* genotype for each strain is shown across the top in dark purple (*plg-1+*), dark green (*plg-1–*), light purple (likely *plg-1+*), light green (likely *plg-1–*), and yellow (ambiguous). The strains which were confirmed by PCR genotyping are also shown. **Alt Text:** Three-panel figure comparing genetic variation between *plg-1+* and *plg-1− C. elegans* strains, including nucleotide diversity across chromosome III, genetic variants per 10 kb window, and *plg-1* haplotype structure across strains.

## MATERIALS AND METHODS

### Worm Culture and Strains

Distinct *C. elegans* genetic backgrounds (*i.e.*, strains) were selected to capture a spectrum of genetic diversity present in the species. We initially screened the 12 strains included in the *Caenorhabditis* Natural Diversity Resource (CaeNDR) divergent set (Crombie et al. 2023). Additional strains were then selected to expand our sample of strains isolated from the Hawaiian Islands (N = 50 strains total) because the highest genetic diversity is found in these strains (Crombie et al. 2019). To support our findings in *C. elegans*, we also included three *C. remanei* strains collected in the eastern United States: QX1539, QX1557, and KRK1. *Caenorhabditis remanei* is an obligate outcrossing species with fixed mate plugging. A complete list of *C. elegans* and *C. remanei* strain information is listed in File S1. Strains were used for PCR genotyping to improve computational benchmarking, while a different set of strains (based on the CaeNDR divergent mapping set) was selected for fecundity analyses to maximize phylogenetic diversity. Three strain pairs were selected from the initial hermaphrodite fecundity analysis for future investigation.

For both species, strains were cultured on 10 cm NGM-agar plates seeded with OP50-1 *Escherichia coli* and maintained at 20°C (Brenner 1974; Kenyon 1988). Prior to all experiments strains were cleaned and age synchronized using hypochlorite treatment (Kenyon 1988). Male-rich *C. elegans* strains were maintained by transferring an additional 30-40 young adult males during the transfer of worms to plates with fresh bacteria.

### *plg-1* Genotyping

We used polymerase chain reaction (PCR) to test for the presence or absence of *Cer1* insertion in *plg-1* using the primers developed in Palopoli *et al*. (2008). Genotype confirmation was performed for the 72 strains listed in File S1. Genomic DNA was purified by isolating two adult worms in 4 μL of 20 mg/μL Proteinase K in elution buffer. Samples were freeze-cracked in liquid nitrogen three times. They were then incubated at 58°C for 60 minutes followed by 10 minutes at 95°C to inactivate the Proteinase K. PCR products were generated using the One-Taq Quick Load 2X Master Mix with Standard Buffer (NEB #M0486) and 2 μL of purified DNA in accordance with manufacturer instructions. Each sample was run for two reactions. One reaction amplified a functional copy of *plg-1* and produced a 599 bp product (forward primer: CGCATAAAACGTCAGCAGAA; reverse primer: ATTCGGAGTAGTCGGGTCCT). The other reaction amplified a non-functional copy of *plg-1* as indicated by a retrotransposon insertion and produced a 658 bp product (forward primer: TCCACAAAACCTGCTGACTG; reverse primer: ATCCACTCGATTTTCGCAAC). Strain genotype was classified based on having the presence of the correct product size in one reaction and no product in the other. Genotype calls were verified with plate-based observations of a mating plug.

### Computational Inference of *plg-1* Genotype

To expand the genotype information to all 1,952 sequenced strains (ECA2199 was excluded due to an issue noted on CaeNDR), we developed a computational pipeline based on the paired-end reads overlapping the insertion site of *Cer1* into *plg-1* and read depth within and around *Cer1*. For each of the 1,953 strains with sequencing reads available from the CaeNDR 20260625 data release (Crombie et al. 2023) (BioProject: PRJNA13758), we identified paired-end reads that mapped around the *plg-1* insertion site. Reads were included in the analysis if they were part of a properly-paired set (both mapped with a forward-oriented upstream and reverse-oriented downstream read end), the upstream-most coordinate of the forward fragment mapped to between 30 and 1000 bp before the insertion site, the reverse oriented fragment mapped downstream of the insertion site (either within *Cer1* or in *plg-1* downstream of the insertion site), and the insert size with less than 9,865 bp (the size of *Cer1* plus 1 kb). The downstream junction was not considered as highly repetitive sequence made it an unreliable marker of *Cer1* insertion. In addition, we found the mean read coverage of a 10 kb non-repetitive stretch of *plg-1* 2.6 kb upstream of the *Cer1* insertion site and the mean read coverage of the *Cer1* sequence, excluding the first and last 1 kb.

To classify strains, we found the mean coverage of read ends covering each base of the first 100 bp of *Cer1* with a mated end mapping upstream in *plg-1* (within) and the number of reads spanning *Cer1* mapping to both sides of the insertion side (spanning). Strains were classified as “plugging” if “within” coverage was zero and at least one “spanning” read and “nonplugging” if there were no “spanning” reads but a mean “within” coverage of at least one. Strains with non-zero values in both categories were assigned “likely plugging” if the “spanning” value was greater and “likely nonplugging” if the “within” value was greater. In cases where both values were zero, the strain was marked as “ambiguous. *Cer1* genomic copy number was inferred by rounding the ratio of mean *Cer1* coverage to mean *plg-1* coverage. Computational genotype classifications and summary statistics are given in File S2.

### Inferring Evolutionary History

We calculated nucleotide diversity (π) at the strain level (Files S3-S4) and isotype level (Files S5-S6). Additionally, we calculated π for isotypes excluding those from the Hawaiian Islands (Files S7-S8). For each analysis, strains/isotypes were classified as having a functional copy of *plg-1*, a non-functional copy of *plg-1*, or being computationally ambiguous. Genomic data (hard-filter VCF) were accessed from the CaeNDR 20250625 data release and π was calculated in 10 kb sliding windows using vcftools (Danecek et al. 2011). Genome-wide and chromosome III diversity results were analyzed in the R Statistical Language (Team 2020). We overlaid *plg-1* genotype on the phylogenetic tree of all 1,953 *C. elegans* (CaeNDR 20250625 data release) using the package phytools (Revell 2024) in R version 4.5.3 (R Core Team 2020).

### Fecundity Assays

We assayed the overall reproductive success of 17 *C. elegans* hermaphrodites with a functional (N = 11 strains) or non-functional (N = 6 strains) copy of *plg-1*. These strains included the 12 in the CaeNDR divergent set along with one additional strain with a functional copy of *plg-1* and four additional strains with a non-functional copy of *plg-1*. Individual hermaphrodites were isolated during late larval stage 4 (L4) onto small NGM-agar plates (35 mm diameter) seeded with 20 μL *E. coli* OP50-1. After 24 hours, each hermaphrodite was transferred to a new small plate to continue laying eggs. Transfers continued every 24 hours until day 7 of adulthood when progeny production had ceased. The total number of progeny were counted as a measure of each hermaphrodite’s reproductive success (File S9). The late L4 progeny were screened for the presence of males as an estimate of the non-disjunction rate for each strain. Three independent assays were conducted for each strain for a total of at least 30 hermaphrodites examined, except for JU346 (N = 29). Fecundity data were analyzed using phylogenetically independent orthonormal contrasts to account for phylogenetic relationships with multiple observations per branch tip following Felsenstein (Felsenstein 2008) and using the R package ape (Paradis and Schliep 2018). Data were censored if a hermaphrodite bagged, desiccated on the petri dish wall, or died before progeny production had ceased. The full strain phylogeny was used to represent the strongest inference of node relationships and tips were dropped to represent the 17 strains of interest using phytools (Revell 2024) in R (R Core Team 2020). Differences in the frequency of males between *plg-1* genotypes was analyzed using a proportions test.

From this group, we selected three pairs of closely related strains with alternative *plg-1* genotypes to represent the different plugging states in an otherwise similar genetic background. Specifically, we selected sister strains ED3017 and MY920, and EG4725 and NIC266, which have functional and non-functional *plg-1* genes, respectively. Additionally, we selected the closely related, but paraphyletic, pair JU346 and JT11398 because no other alternative genotype sister strains were in this group of 17 strains.

We assayed the reproductive success of males from each of the strain pairs when mated to a common pseudo-female background, JK574 (*fog-2*(q71) V). In this strain, hermaphrodite self-sperm production is stopped, turning hermaphrodites into functional females (Schedl and Kimble 1988). This mutation is in a N2 lab-adapted genetic background that is *plg-1–*. We isolated individual JK574 pseudo-females during late L4 onto small NGM-agar plates (35 mm diameter) seeded with 10 μL *E. coli* OP50-1 and maintained them as virgins until the start of the assay. A single male was added to each plate and allowed to mate with the female for 24 hours. After the mating period, males were removed, and females were transferred to a new small plate to continue laying eggs. Transfers continued every 24 hours until day 7 of adulthood when progeny production had ceased. The total number of progeny were counted as a measure of each male’s early adulthood reproductive success (File S10). Strain pairs were assayed at the same time with three independent assays conducted for each pair. At least 25 mated females were examined per strain. Within each strain pair fecundity data were analyzed in R (R Core Team 2020), using a generalized linear model that fit the effect of genotype and replicate on fecundity using a normal distribution. Model distribution was selected using the AIC and log-likelihood output from the R package fitdistrplus (Delignette-Muller and Dutang 2015). The resulting p-values were adjusted for multiple testing (Holm and Sture 1979). All comparisons were in the same direction, so we then pooled strains for more power and ran a generalized linear model that fit the effect of genotype and strain pair on fecundity using a normal distribution. Because the phylogeny of these six strains is unrooted, the multcomp package (Hothorn et al. 2008) was used to perform a Tukey planned comparison test between all strain pairs to determine the effect of phylogenetic groupings.

### Sperm Counts

To count the number of sperm cells transferred in a single mating, individual JK574 pseudo-females were isolated during late L4 onto small NGM-agar plates (35 mm diameter) seeded with 10 μL *E. coli* OP50-1 and maintained as virgins for 24 hours. To identify sperm transferred, we stained males with 10 μM MitoTracker Red CMXRos (Invitrogen M7512) on medium NGM-agar plates (60 mm diameter) for approximately 24 hours. A single stained male was then added to each plate. Males were removed after a mating was observed and left for no more than 4 hours after being placed on a plate. Females were mounted on 1% agarose pads with 400 μM sodium azide. Sperm was counted in real time to allow for accurately moving through the z-stack using a Nikon Ti2 inverted microscope (20× DIC microscope objective, 150 ms fluorescence exposure time). Each ejaculate was counted twice to ensure accuracy (File S11). Sperm count data were analyzed using R (R Core Team 2020). A t-test was performed to determine if mean number of sperm transferred differed between strains within a pair. Additionally, we performed a linear model that fit the effect of male genotype and sperm count.

### Male Signaling Assay

To test if mating plugs act as a male competitive signal, we performed two sequential matings using a full factorial design of *plg-1* genotypes for each of the strain pairs. Thus, for each strain pair there were four mating combinations of first and second male genotypes. Individual JK574 pseudo-females were isolated during late L4 onto small NGM-agar plates (35 mm diameter) seeded with 10 μL *E. coli* OP50-1 and maintained as virgins until the start of the assay. A single male was added to each plate and allowed to mate with the female for 24 hours. After 24 hours, the plate was screened for the presence of embryos to ensure that mating had occurred with the first male (File S12). Those females were then moved to a new plate, and a second male was added. To identify sperm transferred from the second male, we stained males with 10 μM MitoTracker Red CMXRos on medium NGM-agar plates (60 mm diameter) for approximately 20 hours. After a 24-hour mating period with the second, stained male, females were mounted on 1% agarose pads with 400 μM sodium azide. Females were scored in real time using a Nikon Ti2 inverted microscope (20× DIC microscope objective, 150 ms fluorescence exposure time) for the presence of second male sperm in the spermathecae, uterus, and mating plug. Three independent assays were conducted for each strain pair (File S13). We repeated these assays in *C. remanei*. Since mate plugging is a fixed trait, sequential matings were performed within a strain (KRK1, QX1539, and QX1557) and thus there was only a single mating combination. All first males mated represented by the presence of eggs on all plates. Again, three independent assays were conducted for each strain (File S14).

To test if mating plugs delay a second male from mating, we repeated the sequential mating, factorial design in *C. elegans*. However, we reduced the mating time to four hours. In our observations of single male matings (see “Sperm Counts”) we observed that males did not mate with a female more than once during this length of time. Therefore, we hypothesized that a reduced frequency of second male matings corresponded to a delay in time to first mating. Again, individual JK574 pseudo-females were isolated during late L4 onto small NGM-agar plates (35 mm diameter) seeded with 10 μL *E. coli* OP50-1 and maintained as virgins until the start of the assay. A single male was added to each plate and allowed to mate with the female for 4 hours after which the male was removed. Females were left to lay eggs overnight (File S15). A second MitoTracker stained male was added to a plate if eggs were present. After a four-hour mating period, females were mount on 1% agarose pads with 400 μM sodium azide. Females were scored in real time using a Nikon Ti2 inverted microscope (20× DIC microscope objective, 150 ms fluorescence exposure time) for the presence of second male sperm. Two independent assays were conducted for each strain pair (File S16).

Sperm presence data were analyzed using R (R Core Team 2020). A proportions test was performed to determine if the observed number of second male matings deviated from the number of first male matings for each sequential mating combination. Additionally, within each strain pair, we performed generalized linear models that fit the effect of first male genotype and replicate on second male sperm presence. Because we found no effect of first male genotype across all strain pairs, we pooled all strains together for greater power. We performed a generalized linear model that fit the effect of first male genotype, second male genotype, and replicate on second male sperm presence. We also examined the effect of first male genotype of the presence of second male sperm in the spermathecae, uterus, and mating plug independently.

### Plug Area Measurements Over Time

Individual females were isolated during late L4 onto small NGM-agar plates (35 mm diameter) seeded with 10 μL *E. coli* OP50-1 and maintained as virgins until the start of the assay. A single plugging male was added to each plate and allowed to mate with the female for 24 hours. After 24 hours, the plate was screened for the presence of embryos to ensure that mating had occurred. The male was then removed and females were imaged 24-, 48-, and 72-hours post-mating period. This experimental procedure was done in *C. elegans* using JK574 pseudo-females mated with either ED3017, EG4725, or JU346 males. In *C. remanei*, females of KRK1, QX1539, and QX1557 were mated with males from the same strain.

Mating plug size was quantified in *C. elegans* and *C. remanei* using Nikon NIS-Elements software (Nikon Instruments Inc., Tokyo, Japan). Images were captured using a Nikon Ti2 inverted microscope equipped with a 20× DIC microscope objective and 43 ms exposure time. For each image, the mating plug was outlined and measured using the Polygonal ROI tool in NIS-Elements. Each traced ROI was converted to a binary layer, and only the reference binary layer was displayed during subsequent tracing to prevent duplicate or overlapping measurements. The boundary of each mating plug was delineated manually, and area (μm^2^) was obtained using the ROI Statistics tool.

Each plug was measured six times in total – three independent measurements by each of two trained analysts – to minimize observer bias and assess measurement repeatability. A total of 276 plugs were analyzed across the three *C. elegans* strains, and 606 plugs were analyzed across the three *C. remanei* strains (File S17). The number of individuals analyzed per strain and per day varied depending on sample availability. The change in plug area over time was analyzed for each species using a linear model that fit time and the interactions between strain and observer. Potential species effects were also modeled in a general linear framework. All analyses were done using R (R Core Team 2020).

## RESULTS

### *plg-1* Loss is Correlated with Globalization in *C. elegans*

We expanded the genotyping of Palopoli *et al*. (Palopoli et al. 2008) to capture the increase in *C. elegans* sampling (Figure S1). Specifically, we focused on genotyping strains from the Hawaiian Islands, which are likely the ancestral location for *C. elegans* due to the high genetic diversity of strains isolated from this region (Crombie et al. 2019; Rockman et al. 2026). If *plg-1* was lost early in the lineage transition to hermaphroditism, then we would expect few Hawaiian strains to maintain a functional copy of *plg-1*. Our new genotyping included 23 strains from Hawaii, 14 strains from Kauai, six strains from Maui, two strains from Molokai, and five strains from O’ahu. Additionally, we expanded the sampling across the continental United States, Australia, and Europe. In total, we genotyped 72 new strains at the *plg-1* locus (File S1), which brings the total number of PCR confirmed *plg-1* genotypes to 112 strains across datasets. Of these, 96 strains have a functional copy of *plg-1* (*plg-1*+) and 16 have a non-functional copy (*plg-1*–). Importantly, all the strains from the Hawaiian Islands are *plg-1*+, suggesting that *plg-1* was not lost early in the transition to hermaphroditism.

Next, we developed a computational pipeline to classify the *plg-1* genotype in the 1,952 sequenced *C. elegans* strains. Genotypes were inferred based on the counts of reads aligning to *plg-1*, to the retrotransposon *Cer1*, and to the exon 3 insertion overlap (Fig. 1A). The results were validated against the PCR genotyped strains. We found that 86% of strains are *plg-1+* (N = 1,682) and 9% of strains are *plg-1–* (N = 171). Model results for 25 strains were classified as likely-nonplugging, with and average coverage ratio of 38.5 for reads indicating a *Cer1* insertion vs. not. Another 23 strains were classified as likely plugging with a ratio of reads indicating plugging of 19.4 vs. not. The remaining 51 strains were ambiguous (no evidence either way) which we hypothesize is due to lower coverage with an average upstream coverage of 18 reads per bp, less than half of the 44 reads per bp of the remaining strains. Of the 75 genotypes strains, 70 matched the computation plugging or nonplugging calls. Two “likely” calls were confirmed by PCR (one of each) and the remaining three indicated ambiguous strains as *plg-1+*. Genetically similar *C. elegans* strains can be grouped into common isotypes (Andersen et al. 2012). Of the 684 isotypes represented 86% are *plg-1+* (N = 585) and 10% are *plg-1–* (N = 65), replicating the strain-level data. Within an isotype all strains have the same *plg-1* genotype, indicating that *plg-1* polymorphism is at the level of genetic background rather than individual populations.

Mapping genotype information to the strain phylogeny again supports that *plg-1* was not lost in the mating system transition because no strains isolated from the Hawaiian Islands were classified as *plg-1–* (Fig. S1-S2). Instead, the phylogeny supports the conclusions in Palopoli *et al*. (2008) that *plg-1* has been lost during globalization. Specifically, the majority of *plg-1–* strains form monophyletic clades that are sampled from specific regions, such as France, Germany, the Netherlands, and Australia (Fig. S2). These phylogenetic relationships suggest a history of retrotransposon insertion into *plg-1* occurs during population bottleneck events that accompany globalization followed by local population expansion and diversification.

To test this globalization hypothesis, we measured nucleotide diversity (π) in strains grouped by each *plg-1* genotype. If *plg-1* loss is correlated with bottleneck events, then we would expect to see a lower genomic diversity both genome-wide and on chromosome III, which contains *plg-1*, in *plg-1–* strains than in *plg-1+* strains. Conversely, if *plg-1* were selected against in hermaphrodites due to antagonistic function, then we would not expect a correlation between genotype at this locus and genome-wide patterns of diversity, while local diversity at the *plg-1* locus would decrease. Genome-wide the mean nucleotide diversity was an order of magnitude larger in *plg-1+* strains (π = 0.002) compared to *plg-1–* strains (π = 0.0006). Both the median (MW-test: W = 88923560, p < 0.001) and distribution of π values (KS-test: D = 0.67, p < 0.001) were significantly different between genotype groups, even though the maximum diversity was the same (π = 0.02). Nucleotide diversity on chromosome III recapitulated these patterns (Fig. 1). The median nucleotide diversity on chromosome III in *plg-1+* strains (π = 0.002) was significantly larger (MW-test: W = 1701594, p < 0.001) than that in *plg-1–* strains (π = 0.0005). This difference was driven by the complete lack of nucleotide diversity in the chromosome center of *plg-1–* strains, overlapping with the *plg-1* locus (Fig. 1A). Both genotype groups show the characteristic hyperdivergent chromosome arms (Lee et al. 2021; Moya et al. 2025). The counts of variants were also significantly higher in *plg-1+* strains (KS-test: D = 0.90, p < 0.001; Fig. 1B). Further, there is a notable homogeneity surrounding the *plg-*1 locus in strains identified as “likely-nonplugging” and “nonplugging” (Fig. 1C). Both the genome-wide and chromosome III diversity patterns were recapitulated at the isotype level (Tables S1-S2). To confirm that *plg-1* is not maintained simply because the *Cer1* element was lost, we also quantified the number of *Cer1* element copies between genotype groups. We found that *Cer1* copy number ranged from 0-16 with 288 *plg-1+* strains inferred to lack any *Cer1* elements. The mean number of *Cer1* elements did not significantly differ between *plg-1+* and *plg-1* strains (MW-test: W = 138948, p = 0.5). Taken together, these data indicate a genome-wide loss of nucleotide diversity in *plg-1–* strains and support that the loss of *plg-1* function is correlated with population bottlenecks during globalization rather than selection against gene function. Instead, they suggest that *plg-1* is maintained despite it being a male-specific gene in an almost entirely hermaphroditic lineage.

### *plg-1* is Maintained in *C. elegans* by Selection on Males

A possible mechanism maintaining *plg-1* is linkage to a hermaphrodite beneficial gene. Here, purifying selection would not effectively remove *plg-1* from the genome if the benefit of the haplotype exceeded any potential costs of mate plugging when males occurred. Thus, hermaphrodites with a functional *plg-1* would have higher fitness than those without, representing the beneficial haplotype rather than a benefit of *plg-1* itself. To test this hypothesis, we measured the total reproductive success of hermaphrodites from 11 *plg-1*+ strains and six *plg-1*– strains (Fig. 2A). The mean fecundity of *plg-1–* hermaphrodites (x ± SE = 270 ± 3.5, N = 258) trended higher than that of *plg-1*+ hermaphrodites (x ± SE = 245 ± 3.8, N = 391) (Fig. 2B). However, this trend was not significant when accounting for phylogenetic relationships (PGLS: F_1,15_ = 1.5, p = 0.24). To provide additional power, we analyzed the hermaphrodite fecundity data generated by Zhang et al. (Zhang et al. 2021). This dataset quantified lifetime fecundity in 120 strains of which 110 were non-overlapping with our dataset. Again, the mean fecundity of *plg-1–* hermaphrodites (x ± SE = 252 ± 2.8) trended higher than that of *plg-1*+ hermaphrodites (x ± SE = 230 ± 0.46) but was not significantly different when accounting for phylogenetic relationships (PGLS: p = 0.96). Together, these results indicate that *plg-1* does not exist on a reproductive beneficial hermaphroditic haplotype and thus linked selection is not maintaining *plg-1* in the genome.

**Figure 2.**
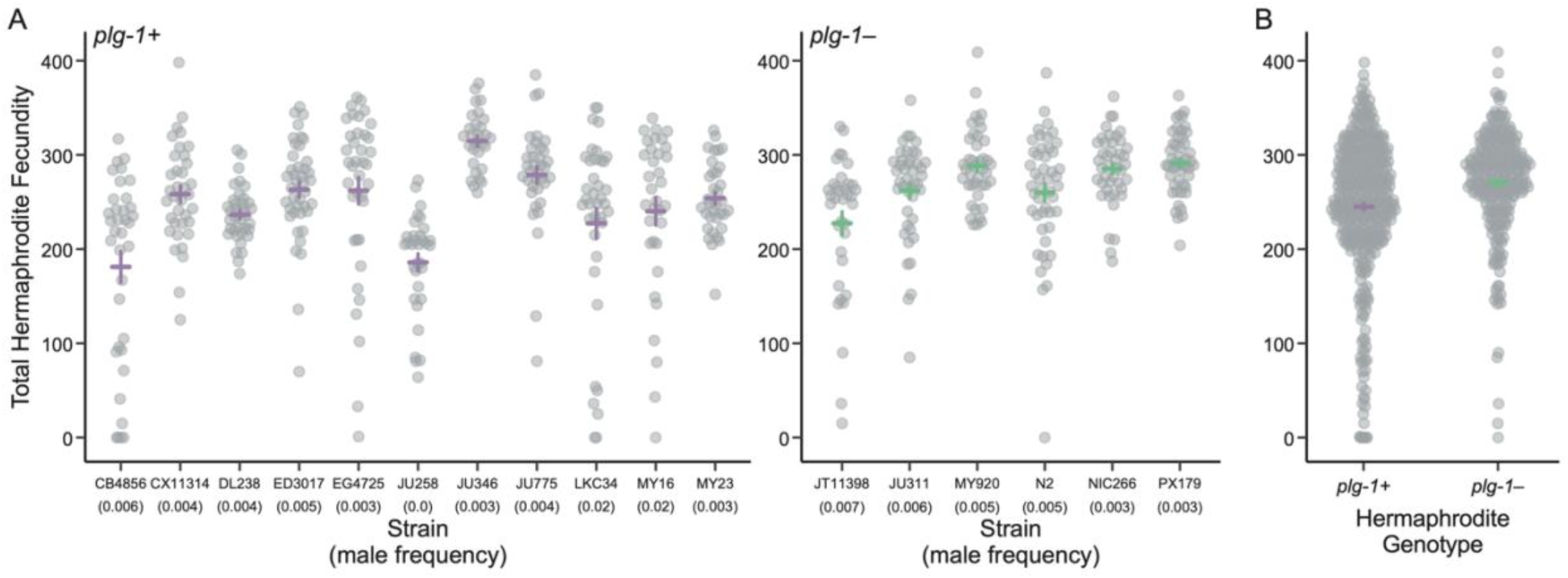
The lifetime fecundity for self-fertilizing *C. elegans* hermaphrodites. **A)** Total fecundity for each of the 11 *plg-1+* and six *plg-1–* strains analyzed. Each point represents the total number of viable offspring for an individual hermaphrodite. At least 33 individual hermaphrodites were examined for each strain. The mean and standard error for the strain is given by the purple (*plg-1+*) and green (*plg-1–*) crosses. The frequency of males observed in each strain due to non-disjunction events are given in parentheses. **B)** Total fecundity grouped by hermaphrodite *plg-1* genotype. Again, each point represents the fecundity of an individual hermaphrodite. The mean and standard error for the strain is given by the purple (*plg-1+*) and green (*plg-1–*) crosses. **Alt Text:** Two-panel figure comparing lifetime fecundity of self-fertilizing *C. elegans* hermaphrodites across eleven *plg-1+* and six *plg-1−* strains, including individual strain fecundity and genotype-level distributions.

Alternatively, *plg-1* could be maintained simply due to differences in male frequency between strains genotypes. Because *C. elegans* has a XX hermaphrodite and XO male sex determination system, males arise in the population through X chromosome non-disjunction events. This explanation would suggest a positive correlation between the maintenance of *plg-1* and non-disjunction rate leading to male production. We quantified the presence of males resulting from non-disjunction during self-fertilization in the same 11 *plg-1*+ strains and six *plg-1*– strains. Male frequency ranged from 0% in JU258 to 2% in both LKC34 and MY16 (Fig. 2A). The proportion of male progeny did not differ significantly between genotypes (χ^2^ = 0.17, df = 1, p = 0.68). Thus, *plg-1* is not maintained through increased male frequency in *plg-1+* populations. Rather, the maintenance of *plg-1* must be due to a selective advantage of its function during reproduction.

### Mating Plugs Do Not Prevent or Delay Future Reproductive Events

Next, we tested aspects of the paternity assurance hypothesis to determine if the presence of a mating plug either chemically or physically prevents or delays a second male from mating. We sequentially mated feminized *C. elegans* hermaphrodites (*i.e.*, hermaphrodites that cannot produce self-sperm and therefore are functionally female) with *plg-1*+ and *plg-1*– males from three strain pairs using a full factorial design. We then quantified the presence of fluorescently dyed sperm from the second male across three anatomical regions: spermathecae, uterus, and mating plug (Fig. S2A). If mating plugs are a male competitive signal, then we would expect to see fewer sperm from the second male in the female reproductive tract if a mating plug was present.

When given sufficient time to mate more than once, the presence of second male sperm ranged from 81% to 94% (Fig. 3A; Fig. S2B-C). Fewer second males mated than first males only when both males were *plg-1–* (χ^2^ = 22.5, df = 1, p < 0.001). We found no difference in first and second male mating frequency when the first male was *plg-1+* regardless of second male genotype (p = 0.2; p = 0.7). Moreover, we found no effect of first male genotype on second male sperm presence within any region of the female reproductive tract for each strain pair (LM: F_4,619_ = 3.3, t = 0.73, p = 0.46). The lack of effect did not change when analyzing sperm presence separately for each anatomical region (LM*_spermathecae_*: F_1,622_ = 0.58, p = 0.45; LM*_uterus_*: F_1,622_ = 0.81, p = 0.37; LM*_plug_*: F_1,622_ = 0.58, p = 0.44). Thus, mating plugs do not impede a male from mating or limit a female’s access to sperm. However, we detected a significant effect of second male genotype on sperm presence such that when the second male produced a mating plug a higher frequency of females were observed with second male sperm (LM: F_4,619_ = 3.3, t = 3.5, p < 0.001).

**Figure 3.**
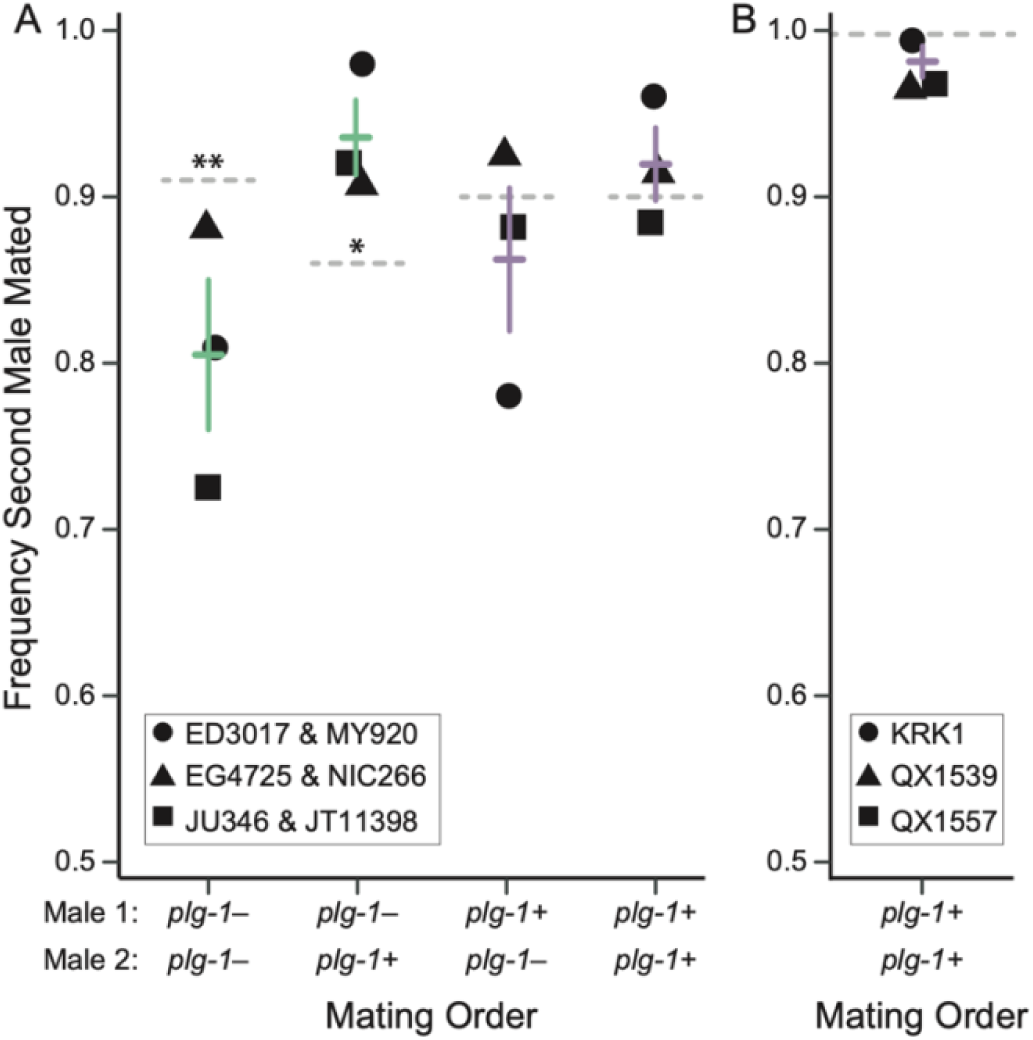
The frequency that sperm from the second mating male in a sequential mating is observed within the female reproductive tract. **A)** Factorial crosses in *C. elegans* for each of the three strain pairs examined are given by the different shaped points. Each point represents the frequency observed in at least 45 mated females. The mean and standard error across strains is shown and colored based on the first male genotype (green is *plg-1–* and purple is *plg-1+*). The dashed line represents the mean mating frequency of the first males. Asterisks denote a significant difference in the proportion of first and second males mating (*p < 0.05, **p < 0.001). **B)** Sequential cross within three *C. remanei* strains. Each point represents the frequency observed in at least 35 scored females from each strain with the shapes representing the different strains. The mean and standard error across strains is shown. The dashed line represents the 100% mating frequency of the first males. **Alt Text:** Two-panel figure comparing the frequency of successful second-male mating following sequential matings involving different male strains and mating orders in *C. elegans* and *C. remanei*.

We repeated these assays in *C. remanei*, a gonochoristic species where mate plugging is a fixed trait. In all three strains examined the second male again mated more frequently than expected under random mating (χ^2^ = 98.23, df = 1, p < 0.001). The frequency of second male matings was higher than in *C. elegans* and ranged from 97% to 100% (Fig. 3B). Again, we found no effect of first male genotype on second male sperm transfer to any region of the female reproductive tract (LM: F_1,105_ = 0.50, p = 0.61), providing more statistically conclusive evidence than previous work (Timmermeyer et al. 2010).

To test if mating plugs delay the time to mating, we repeated these assays in *C. elegans* using a shorter mating time where males were unlikely to mate more than once. Fewer males overall were observed to mate (Fig. S4). We found a marginally non-significant effect of first male genotype on second male sperm presence (LM: F_3,104_ = 1.4, p = 0.05), such that females with a mating plug were mated more frequently than by a second male. In fact, second male mating frequency was significantly lower when the first male was *plg-1–* (χ^2^ = 8.58, df = 2, p = 0.01). These results demonstrate that if a male is deterred from mating a non-virgin female, the mating plug itself was likely not the cause. Together, these data do not support that mating plugs in *Caenorhabditis* nematodes act as a strong male competitive deterrence and provide no evidence that they limit female access to future matings.

### Females with a Mating Plug Have Higher Fecundity

Given the lack of evidence for the paternity hypothesis, we then tested if instead mate plugging provides a female benefit. *Caenorhabditis elegans* pseudo-females were mated with a single male from each of the three strain pairs and the progeny produced was quantified across days of adulthood. The total progeny produced ranged from 184 offspring (N = 39) in JT11398 to 343 offspring (N = 42) in EG4725 (Fig. 4). In females mated with *plg-1*+ males, reproduction peaked on day 3 of adulthood and was consistently higher than females mated with *plg-1–* males (Fig. 4A). We found a significant effect of genotype (GLM: t-value = 3.17, p < 0.01) and day (GLM: t-value = −13.1, p < 0.001) on female fecundity, but not the interaction between them (p = 0.7). We detected a significant effect of strain pair such that pair JU346/JT11398 was significantly different from pair ED3017/MY920 (MPC: z = 3.5, p = 0.001) and pair EG4725/NIC266 (MPC: z = 2.6, p = 0.02), likely driven by the low fecundity observed in JT11398. We found no strain effect between pairs ED3017/MY920 and EG4725/NIC266 (p = 0.6). This difference in progeny laid per day translated to females producing on average 100 more offspring when mated with *plg-1*+ males (x ± SE = 354 ± 1.3, N = 97) than when mated with *plg-1–* males (x ± SE = 246 ± 1.4, N = 107) (Fig. 4B). This difference in total fecundity was significant for the individual pairs JU346/JT11398 (GLM: t-value = 4.7, p_adj_ < 0.001) and ED3017/MY920 (GLM: t-value = 3.5, p_adj_ < 0.01), but only trending in pair EG4725/NIC266 when adjusting for multiple comparisons (p = 0.1).

**Figure 4.**
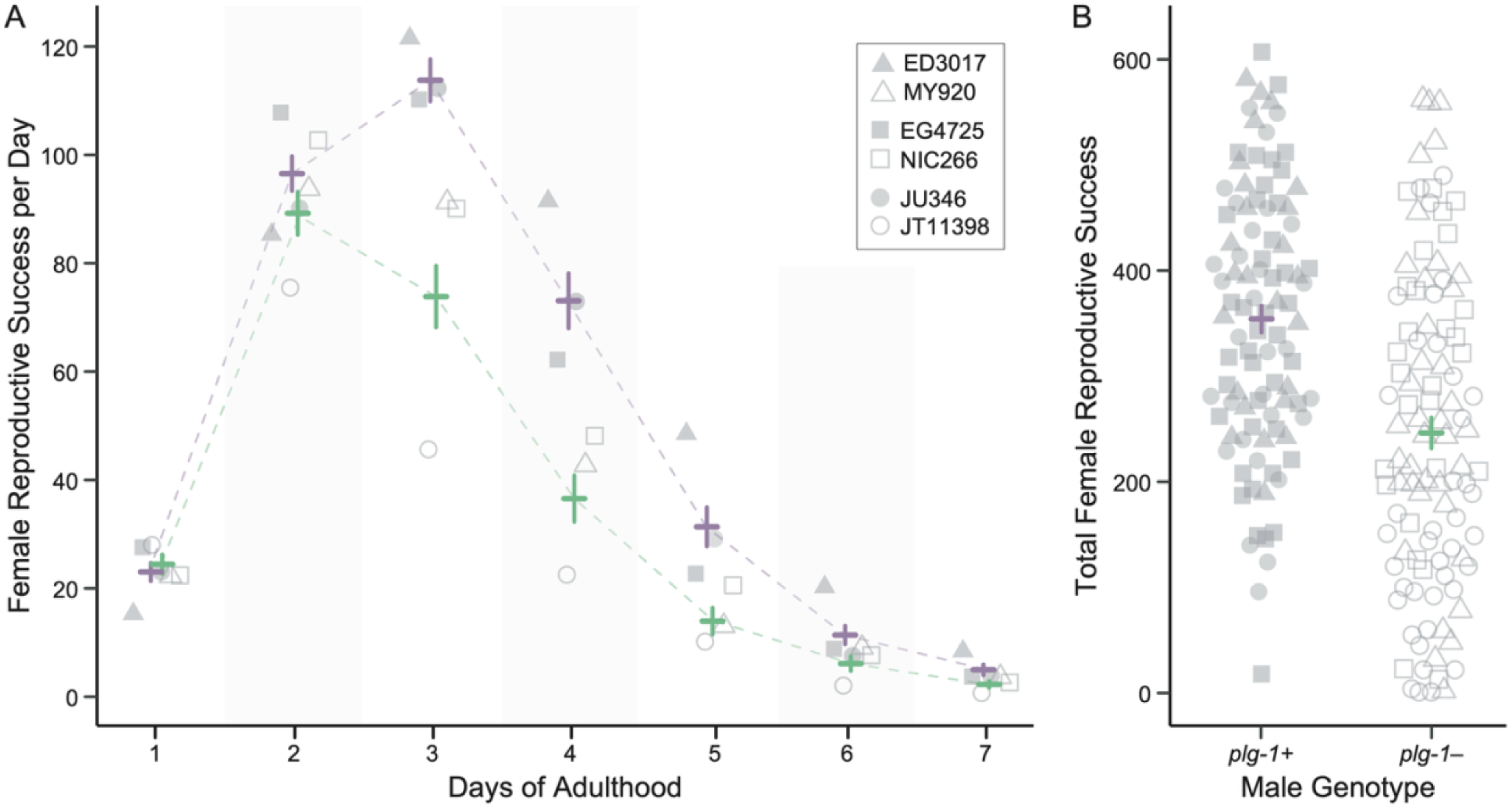
The fecundity of *C. elegans* pseudo-females mated with either *plg-1+* or *plg-1–* males. **A)** Female reproductive success across days of adulthood. Each point represents the mean fecundity of at least 25 mated females. The point shapes represent the male genotype for the sister strain pairs with filled shapes corresponding to the *plg-1+* genotype and the open shapes corresponding to the *plg-1–* genotype. The mean and standard error across strains of the same genotype is shown (green is *plg-1–* and purple is *plg-1+*). **B)** Total female fecundity summed across all of adulthood and grouped by male *plg-1* genotype. The point shapes are the same as in panel A. The mean and standard error is shown. **Alt Text:** Two-panel figure comparing daily and lifetime reproductive success of *C. elegans*pseudo-females mated with *plg-1+* and *plg-1−* males.

We found that the number of sperm transferred in a single mating did not differ significantly between strain pairs of by male genotype (LM: F_1,69_ = 1.03, p = 0.3), suggesting that differences in sperm count likely do not explain the differences in fecundity observed (Fig. S5). Additionally, mean sperm transfer was not consistently higher in *plg-1+* strains. However, the high variance in sperm transferred within and between strains necessitates sample sizes over 2,500 individuals per genotype to have 80% power to detect a significant difference of the effect size quantified. Overall, the presence of a mating plug alone appears to increase female reproductive success by lengthening peak reproduction and total reproductive success.

### Plug Size Does Not Decrease with Time Post-Mating

Mating plugs have been shown to benefit female fitness by acting as a nuptial gift (Perry and Rowe 2008) or preventing sperm loss (Avila et al. 2015). If the mating plug in *Caenorhabditis* nematodes were acting a nutrient resource, then we might expect the size of the plug to decrease post-mating as it was broken down to macromolecular components. Alternatively, if the plug were instead acting as a vulval cover to retain sperm, then we would expect to see no change in plug size over time. To examine these alternative forms of benefits, we mated *C. elegans* pseudo-females and *C. remanei* females with *plg-1*+ males of the respective species and then measured plug area 24-, 48-, and 72-hours post-mating.

In *C. elegans*, plug area was greatest 72-hours post-mating (x ± SE = 2485 ± 13.6um^2^, N = 102) and smallest 48-hours post-mating (x ± SE = 2085 ± 40.2um^2^, N = 60), though sample size was also smallest on this day (Fig. 5A). Overall, we found no significant change in plug area over time, nor did plugs differ between male genotypes (LM: F_6,269_ = 1.10, p = 0.36). In *C. remanei*, plug area increased from 24-hours post-mating (x ± SE = 2625 ± 7.8um^2^, N = 180) to 72-hours post-mating (x ± SE = 3884 ± 11.3um^2^, N = 216) in a significant fashion (Fig. 4B; LM: F_6,591_ = 41.9, p < 0.001). Although, we detected differences in plug area between genotypes, suggesting that expression level could vary across genetic backgrounds (Fig. 5B), each strain independently had a significant increase in area over time (LM_KRK1_: F_1,226_ = 18.2, p < 0.001; LM_QX1539_: F_1,184_ = 35.1, p < 0.001; LM_QX1557_: F_1,184_ = 12.0, p < 0.001). Finally, we examined the species effect and found a significant interaction between species and time (LM: t-value = 3.14, p < 0.01), despite each individual component not contributing significantly to plug area. Together, these results show that plug area is maintained or increased with time post-mating in a species-specific manner.

**Figure 5.**
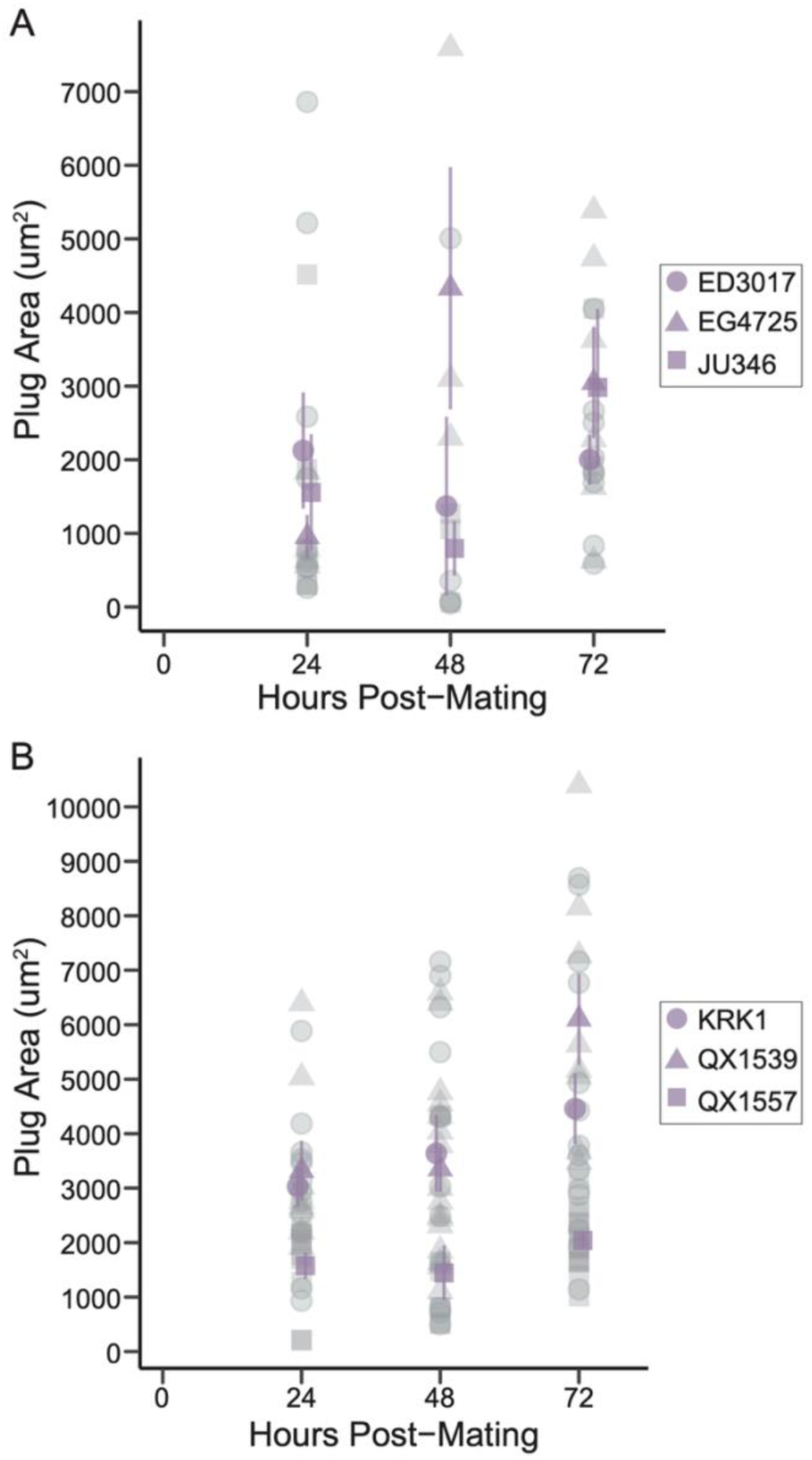
Mating plug area 24-, 48-, and 72-hours post-mating. **A)** Plug area measured on *C. elegans* pseudo-females mated with three different strains of *plg-1+* males (represented by the different shapes). Each point represents the mean area of plug from six measurements (three technical replicate measures from each of two independent observers). The mean and standard error for females mated with each male genotype are shown in purple. **B)** Plug area measured on mated *C. remanei* females from three different strains (represented by the different shapes). Again, each point represents the mean area of plug from six measurements (three technical replicate measures from each of two independent observers). The mean and standard error for females mated with each male genotype are shown in purple. **Alt Text:** Two-panel figure comparing mating plug area at 24, 48, and 72 hours post-mating across different male strains of *C. elegans* and *C. remanei*.

## DISCUSSION

Studying the evolutionary transition to self-fertilizing hermaphroditism provides a comparative framework to probe why and what types of genes with sex-specific functions are lost versus maintained. Here, we examined *plg-1,* a male-expressed gene that underlies mate plugging, though the function it plays in reproduction is not well characterized. The repeated loss of other male-specific genes would suggest genome streamlining to lose *plg-1* during the lineage transition to self-fertilizing hermaphroditism. Instead, we found that *plg-1* has been maintained in at least 86% of *C. elegans* strains, including all the ancestral Hawaiian Island strains. The loss of mate plugging likely occurred several independent times during globalization. We find no evidence that mate plugging prevents, impedes, or delays another male from mating nor does it appear to limit female mating choice. Rather, the mating plug appears to act as a beneficial cover that prolongs female reproductive success post-mating and increases overall fecundity. Together, these results indicate that male-specific expression does not equate to male-specific function and demonstrate the role male-specific genes play during reproduction even when genetic males are not required for reproduction to occur.

Both genetic-diversity estimates and phylogenetic relationships indicate that the loss of *plg-1* is correlated with range expansion out of the Hawaiian Islands. During such globalization genetic diversity decreases due to the coupling of population bottlenecks and serial founder effects (Henn et al. 2012). In *C. elegans*, a single hermaphrodite can found a new population. Thus, if the *Cer1* retrotransposon were to insert into *plg-1* in a single founder, then all subsequent generations would contain the insertion and lack a function copy of *plg-1*. This scenario likely explains the 10% loss of *plg-1* across strains. A loss of the *Cer1* element is not responsible for the maintenance of *plg-1*. Interestingly, although we annotated a median of four *Cer1* copies in *plg-1+* strains, the maximum copy number was four times higher. Previous work in on these elements in a lab-adapted *C. elegans* strain annotated only four *Cer1* elements (Ganko et al. 2001), suggesting that this retrotransposon family could have a more complex evolutionary history in the genetically diverse Hawaiian strains.

Early observations of mate plugging suggested that it was a mutually beneficial reproductive trait (Voss 1981). Although paternity assurance and sexual conflict is seen particularly in some mammalian species {see Mangels.2016}, the broad distribution of mate plugging suggests that reproductive benefits of a mate plugging to their mating partners are likely the predominant function (Perry and Rowe 2008; Avila et al. 2015; Schneider et al. 2016; McDonough-Goldstein et al. 2022; Yun et al. 2024). Supporting this perspective, we find no evidence for the paternity assurance hypothesis and instead show that females with a mating plug have increased fertility. Although our experimental design maximized the effect of sperm limitation by using *C. elegans* pseudo-females, it is directly comparable to and expands on expands on previous work in *C. remanei* that showed that plugged females produced more eggs and offspring than non-plugged females (Timmermeyer et al. 2010). Due to the high variance in sperm transferred both within and across species (also see Murray and Cutter 2011), we find it unlikely that sperm count is the sole driver of this effect. Thus, these strain pairs support the hypothesis that *Caenorhabditis* mating plugs increase female fecundity along with increasing the fecundity of the depositing male and any potential future male mates.

Although we cannot rule out the potential that the mating plug acts as a nutrient gift, we find no evidence that plug area decreases post-mating as would be expected if it were actively broken down into its component macromolecular parts. Additionally, the nutrient content of a mucin protein is not immediately clear. Instead, we hypothesize that the mating plug acts as vulval cover that allows males to still find the vulva opening for copulation but does not allow sperm cells to pass out of the reproductive tract. Previous studies showed that females pressurize the reproductive tract to push out the male spicule after mating and that sperm can also be lost during this process (Barker 1994; Kleemann and Basolo 2007). This sperm loss can lead to females being sperm limited (Kleemann and Basolo 2007). We show two lines of circumstantial evidence that the mating plug retains sperm. First, the higher late-life fecundity of plugged females coupled with visualized sperm within the mating plug suggest that the plug prevents sperm from being pushed out of the uterus during vulval muscle contractions. Second, plug area was maintained post-mating or even increased in size, suggesting the cover itself is important. Together, these results support that at least one function of mating plugs is to prevent sperm loss and represent a promising avenue for future research. Further, our work builds on a body of literature and demonstrates the power of including nematodes to integrate genetics, reproductive fitness, and function on the study of mate plugging.

## Data Availability

The complete list of strains analyzed in this study along with their *plg-1* genotype are given in File S1. Computationally inferred *plg-1* genotype and *Cer1* copy number are given in File S2. Nucleotide diversity statistics are given in Files S3-S8. The hermaphrodite fertility data are available in File S9, and the pseudo-female fertility data are available in File S10. The 24-hour plug signaling data are available in File S11-S12 (*C. elegans*) and File S13-S14 (*C. remanei*). The 4-hour plug signaling data are available in File S15-S16. Plug area data are available in File S17. All scripts are available via the Kasimatis Lab GitHub repository plg-1 (https://github.com/Kasimatis-Lab/plg-1). All *C. elegans* strains are available from CaeNDR, the *Caenorhabditis* Natural Diversity Resource. Strains QX1539 and QX1557 were generously provided by the Andersen lab. Strain KRK1 is available from the Kasimatis lab upon request.

## Author Contributions

KRK devised the project. LJB performed the PCR genotyping, and MS performed the computational genotype inference. LJB and KRK performed the fecundity assays. LJB performed the plug signaling assays with assistance from KRK. LJB and AG quantified plug area over time. KRK analyzed the data. KRK wrote the manuscript with the support of the other authors.

## Acknowledgements

Maham Malik and Joanne Yang assisted with counting the hermaphrodite fecundity assays. Worm strains were provided by CaeNDR, which is funded by NSF Grant 1930382. Some strains were provided by the CGC, which is funded by NIH Office of Research Infrastructure Programs (P40 OD010440). Additionally, we thank the Andersen lab for sharing *C. remanei* strains. We thank Drew Schield for assistance in calculating genetic diversity. We thank Locke Rowe and the Kasimatis lab for their helpful discussions.

---

**Table S1.** Genome-wide diversity summary statistics. Genetic backgrounds are grouped by *plg-1* genotype using either strain-level or isotype-level data. Isotypes were also analyzed excluding all Hawaiian Island genetic backgrounds (N = 262). The mean, minimum, and maximum genetic diversity (π) are given along with the mean number of variants per 100 kb window.

| *plg-1* Genotype | Grouping | Mean π | Min π | Max π | Mean # Variants |
|---|---|---|---|---|---|
| plugging | strains | 0.002 | 0.000000004 | 0.02 | 355 |
|  | isotypes | 0.002 | 0.000000003 | 0.02 | 349 |
|  | isotypes, excluding HW | 0.001 | 0.000000008 | 0.02 | 134 |
| non-plugging | strains | 0.0006 | 0.00000003 | 0.02 | 50 |
|  | isotypes | 0.0007 | 0.00000007 | 0.02 | 50 |

**Table S2.** Diversity summary statistics for chromosome III. Genetic backgrounds are grouped by *plg-1* genotype using either strain-level or isotype-level data. Isotypes were also analyzed excluding all Hawaiian Island genetic backgrounds (N = 262). The mean, minimum, and maximum genetic diversity (π) are given along with the mean number of variants per 100 kb window.

| *plg-1* Genotype | Grouping | Mean π | Min π | Max π | Mean # Variants |
|---|---|---|---|---|---|
| plugging | strains | 0.002 | 0.000001 | 0.01 | 305 |
|  | isotypes | 0.002 | 0.000002 | 0.01 | 299 |
|  | isotypes, excluding HW | 0.001 | 0.0000004 | 0.01 | 123 |
| non-plugging | strains | 0.0005 | 0.0000004 | 0.01 | 48 |
|  | isotypes | 0.0006 | 0.0000004 | 0.01 | 49 |

**Figure S1.** Phylogenetic relationships between *C. elegans* strains. Tip color corresponds to the computationally inferred *plg-1* genotype, where *plg-1*+ strains are dark purple, *plg-1–* strains are dark green, likely *plg-1+* strain are light purple, likely *plg-1–* strains are light green, and ambiguous genotypes are yellow. The phylogeny was generated in the CaeNDR 20250625 data release.

**Alt Text:** Phylogenetic tree showing evolutionary relationships among *C. elegans* strains, with branches colored according to computationally inferred *plg-1* genotype.

**Figure S2.** The geographic distribution of *C. elegans* isotype. Point color corresponds to the computationally inferred *plg-1* genotype, where *plg-1*+ isotypes are purple, *plg-1–* isotypes are green, and ambiguous genotypes are gray.

**Alt Text:** Scatter plot showing the distribution of *C. elegans* isotypes by latitude and longitude, with points colored according to *plg-1* genotype.

**Figure S3.** Representative images of scoring second male sperm presence using MitoTracker Red CMXRos. **A)** A schematic diagram of a female nematode with the spermathecae, uterus, and mating plug labeled. The colored dots represent dyed sperm cells. **B)** A representative female with second male sperm in the spermathecae (S), uterus (U), and mating plug (P). From left to right the images were taking using DIC brightfield light and red fluorescence using a Nikon Ti2 inverted microscope (20× DIC microscope objective, 150 ms fluorescence exposure time). The merged image is shown on the right. This female was mated first with a JU346 (*plg-1+*) male and second with another JU346 male. **C)** Representative images of the factorial crossing design between ED3017 (*plg-1+*) and MY920 (*plg-1–*). The merged brightfield and fluorescence image is shown.

**Alt Text:** Three-panel figure illustrating the detection and localization of fluorescently labeled second-male sperm in female nematodes using reproductive anatomy schematics and representative microscopy images.

**Figure S4.** The frequency that sperm from the second mating male in a sequential mating (4-hour windows) is observed within the female reproductive tract. Factorial crosses in *C. elegans* for each of the three strain pairs examined are given by the different shaped points. The mean and standard error across strains is shown and colored based on the first male genotype (green is *plg-1–* and purple is *plg-1+*). The dashed line represents the mean mating frequency of the first males. Asterisks denote a significant difference in the proportion of first and second males mating (*p < 0.05).

**Alt Text:** Scatter plot comparing the frequency of second-male sperm observed in the female reproductive tract across four sequential mating combinations involving *plg-1+* and *plg-1− C. elegans* males.

**Figure S5.** The number of sperm transferred in a single ejaculate. **A)** Each point represents the count of sperm cells transferred to a female during a single mating event for each strain. The mean and standard error within a strain is shown and colored based on the male genotype (green is *plg-1–* and purple is *plg-1+*). There is no significant difference in mean sperm transferred within strain pairs or across all strains. **B)** The grouped distribution of sperm transferred across all males. The mean number of sperm transferred in *plg-1+* strains (purple line) is six sperm cells more than in *plg-1–* strains (green line), though this is not significant.

**Alt Text:** Two-panel figure comparing the number and distribution of sperm transferred during individual mating events across *plg-1+* and *plg-1− C. elegans* strains.

## Supplemental Files

**File S1. Strain data.** The complete list of strains analyzed in this study along with their *plg-1* genotype.

**File S2: Inferred genotype for 1,953 *C. elegans* strains.** The strain, inferred *plg-1* genotype, inferred number of *Cer1* elements along with their location, *plg-1* coverage, *Cer1* coverage, and orphan reads are given.

**File S3: Genome-wide nucleotide diversity statistics for *C. elegans* strains.** The chromosome, window start (in bp), window end (in bp), number of variants, nucleotide diversity, and *plg-1* genotype are given.

**File S4: Chromosome III nucleotide diversity statistics for *C. elegans* strains.** The chromosome, window start (in bp), window end (in bp), number of variants, nucleotide diversity, and *plg-1* genotype are given.

**File S5: Genome-wide nucleotide diversity statistics for *C. elegans* isotypes.** The chromosome, window start (in bp), window end (in bp), number of variants, nucleotide diversity, and *plg-1* genotype are given.

**File S6: Chromosome III nucleotide diversity statistics for *C. elegans* isotypes.** The chromosome, window start (in bp), window end (in bp), number of variants, nucleotide diversity, and *plg-1* genotype are given.

**File S7: Genome-wide nucleotide diversity statistics for *C. elegans* isotypes excluding those from the Hawaiian Islands.** The chromosome, window start (in bp), window end (in bp), number of variants, nucleotide diversity, and *plg-1* genotype are given.

**File S8: Chromosome III nucleotide diversity statistics for *C. elegans* isotypes excluding those from the Hawaiian Islands.** The chromosome, window start (in bp), window end (in bp), number of variants, nucleotide diversity, and *plg-1* genotype are given.

**File S9: Hermaphrodite total fertility.** The total reproductive success for hermaphrodites from 11 *plg-1+* strains and 6 *plg-1–* strains. Data were censored (1) if a hermaphrodite bagged, desiccated on the petri dish wall, or died before progeny production had ceased.

**File S10: Pseudo-female fecundity.** The reproductive success of *C. elegans* pseudo-females mated with either a *plg-1+* male or a *plg-1–* male for 24 hours. Data were censored (1) if a female bagged or desiccated on the petri dish wall.

**File S11: Sperm cell transfer.** The counts of sperm cells transferred to *C. elegans* pseudo-females from a single mating with either a *plg-1+* male or a *plg-1–* male.

**File S12: Plug signaling null distribution in *C. elegans*.** The frequency of eggs observed on the plate from a first male during sequential factorial crosses in three *C. elegans* sister strain pairs. Matings lasted for 24 hours.

**File S13: Plug signaling in *C. elegans*.** The frequency of observed sperm transferred from a second male during sequential factorial crosses in three *C. elegans* sister strain pairs. Matings lasted for 24 hours.

**File S14: Plug signaling in *C. remanei*.** The frequency of observed sperm transferred from a second male during sequential crosses in three *C. remanei* strains. All first males mated.

**File S15: Delay in mating null distribution in *C. elegans*.** The frequency of eggs observed on the plate from a first male during sequential factorial crosses in three *C. elegans* sister strain pairs. Matings lasted for four hours.

**File S16: Delay in mating in *C. elegans*.** The frequency of observed sperm transferred from a second male during sequential factorial crosses in three *C. elegans* sister strain pairs. Matings lasted for four hours.

**File S17: Plug area post-mating.** The change in plug area 24-, 48-, and 72-hours post-mating in three *C. elegans* strains and three *C. remanei* strains.

