## Supplementary figures and images for "A Male Expressed Copulatory Plug Increases Female Fertility in *Caenorhabditis* Nematodes"

### Figure S1

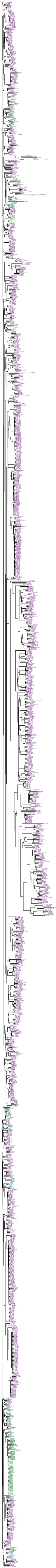

### Figure S2

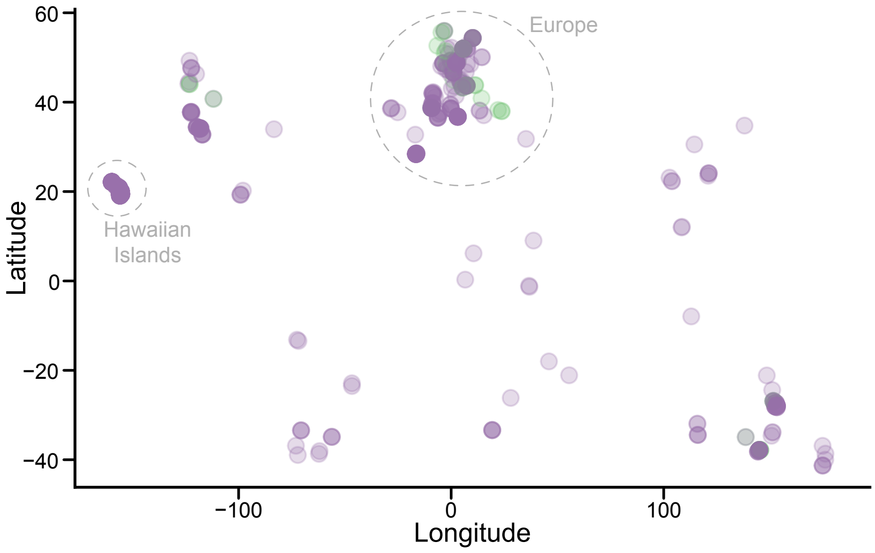

Europe
Hawaiian Islands
Latitude
Longitude
60
40
20
0
−20
−40
−100
0
100

### Figure S5

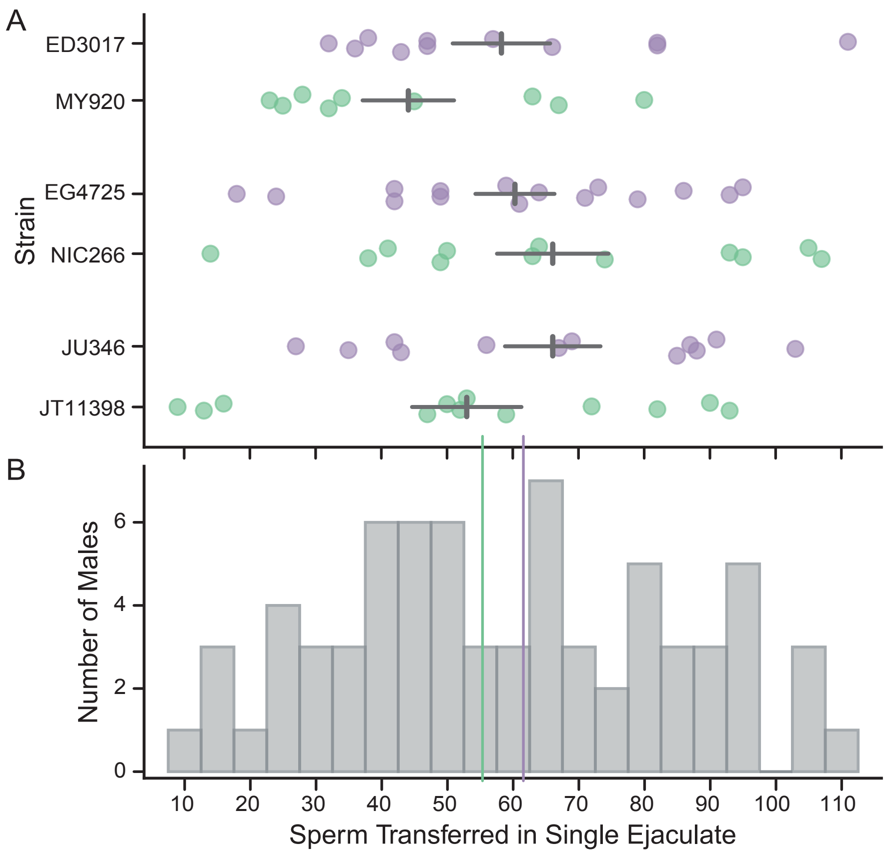

A
B
Strain
ED3017
MY920
EG4725
NIC266
JU346
JT11398
Number of Males
0
2
4
6
10
20
30
40
50
60
70
80
90
100
110
Sperm Transferred in Single Ejaculate
