## Supplementary material for "A Male Expressed Copulatory Plug Increases Female Fertility in *Caenorhabditis* Nematodes": Figure S3

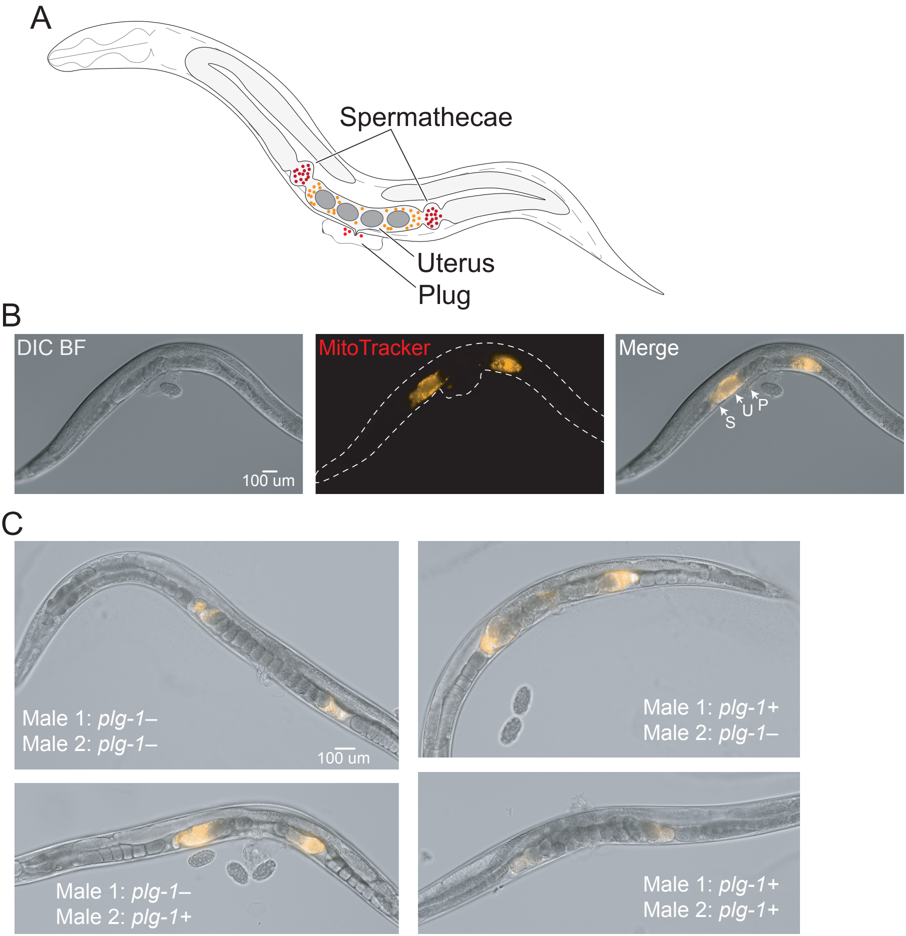
A
Spermathecae
Uterus
Plug
B
DIC BF
100 um
MitoTracker
Merge
S
U
P
C
Male 1: plg-1–
Male 2: plg-1–
100 um
Male 1: plg-1+
Male 2: plg-1–
Male 1: plg-1–
Male 2: plg-1+
Male 1: plg-1+
Male 2: plg-1+
