## Supplementary material for "A Male Expressed Copulatory Plug Increases Female Fertility in *Caenorhabditis* Nematodes": Figure S4

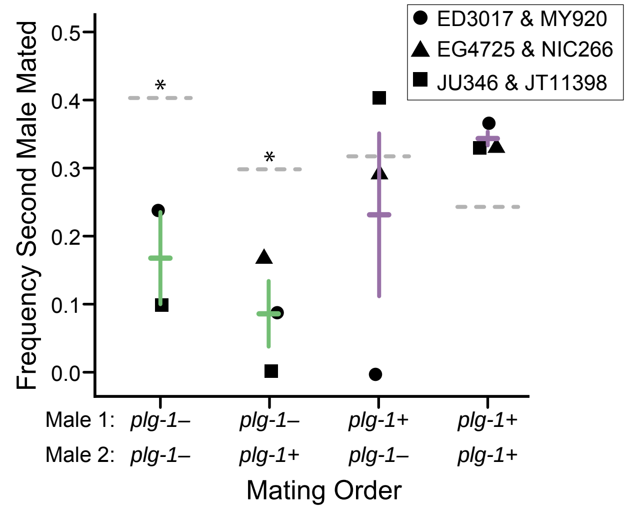
ED3017 & MY920
EG4725 & NIC266
JU346 & JT11398
Frequency Second Male Mated
0.5
0.4
0.3
0.2
0.1
0.0
*
*
Male 1: plg-1– plg-1– plg-1+ plg-1+
Male 2: plg-1– plg-1+ plg-1– plg-1+
Mating Order
